# Serotonin Promotes Vesicular Association and Fusion by Modifying Lipid Bilayers

**DOI:** 10.1101/2024.01.20.576155

**Authors:** Debsankar Saha Roy, Ankur Gupta, Vicky Vishvakarma, Pawel Krupa, Mai Suan Li, Sudipta Maiti

**Affiliations:** Dept. of Chemical Sciences, Tata Institute of Fundamental Research, Homi Bhabha Road, Colaba, Mumbai 400005, India; Institute of Physics, Polish Academy of Sciences, Warsaw 02-668, Poland; Institute for Computational Science and Technology, Quang Trung Software City, Tan Chanh Hiep Ward, District 12, Ho Chi Minh City, Vietnam

**Keywords:** Exocytosis, Vesicle fusion, Neurotransmitter-lipid interaction, Serotonin, membrane mechanics, small molecule membrane interaction, lipid order

## Abstract

The primary event in chemical neurotransmission involves the fusion of a membrane-limited vesicle at the plasma membrane and the subsequent release of its chemical neurotransmitter cargo. The cargo itself is not known to have any effect on the fusion event. However, amphiphilic monoamine neurotransmitters (e.g. serotonin and dopamine) are known to strongly interact with lipid bilayers and to affect their mechanical properties, which can in principle impact membrane-mediated processes. Here we probe whether serotonin can enhance the association and fusion of artificial lipid vesicles *in vitro*. We employ Fluorescence Correlation Spectroscopy and Total Internal Reflection Fluorescence microscopy to measure the attachment and fusion of vesicles whose lipid compositions mimic the major lipid components of synaptic vesicles. We find that association between vesicles and supported lipid bilayers are strongly enhanced in a serotonin dose-dependent manner, and this drives an increase in the rate of spontaneous fusion. Molecular dynamics simulations and fluorescence spectroscopy data show that serotonin insertion increases the water content of the hydrophobic part of the bilayer. This suggests that the enhanced membrane association is likely driven by an energetically favourable drying transition. Other monoamines such as dopamine and norepinephrine, but not other related species such as tryptophan, show similar effects on membrane association. Our results reveal a lipid bilayer-mediated mechanism by which monoamines can themselves modulate vesicle fusion, potentially adding to the control toolbox for the tightly regulated process of neurotransmission *in vivo*.

**TOC graphics:** 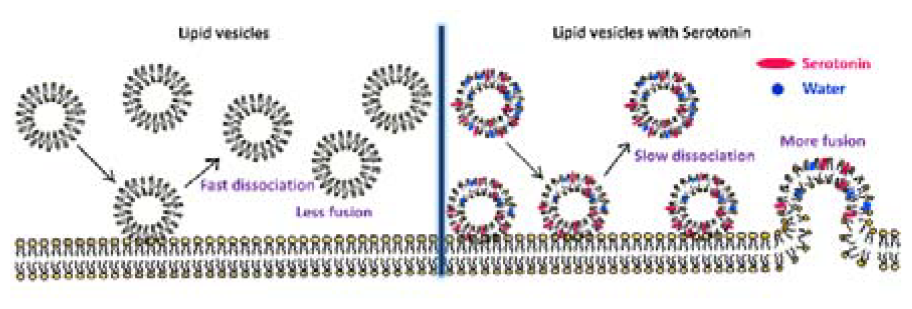

## 1. Introduction

Chemical neurotransmission occurs through the process of regulated exocytosis of neurotransmitter-containing lipid vesicles at the plasma membrane.^1^ It has been reported that the neurotransmitter concentration of the vesicles can be correlated with the exocytosis probability.^2^ However, there is no known physical mechanism by which the intravesicular neurotransmitters, before their release, can exert any control on the vesicle fusion process in the pre-synaptic cell. However, recent studies show that the monoamine neurotransmitters interact with lipid bilayers in addition to their cognate receptors.^3–6^ This interaction can also change the mechanical properties of lipid bilayers.^7–10^ Specifically, a few mM of serotonin was found to decrease the lipid membrane order significantly, as reported by AFM force indentation and ^2^H-NMR measurements.^8^ Also, the ordered domains of a ternary mixture of lipids get restructured by serotonin ^7^, an effect that is also mimicked by serotonergic drug molecules^11^. The change in mechanical properties induced by monoamines can in principle modulate membrane-mediated processes. It is known that fusion pore opening and expansion are energetically costly. It involves significant rearrangement of lipid membranes, so mechanical properties such as bending rigidity and curvature of the membrane should also affect the process of membrane fusion.^12–14^ We do find serotonin-induced changes in the mechanical properties of lipids in model membranes mimicking the vesicular composition^10^. So this can provide a potential mechanism for modulating the exocytosis process.

The existence of such modulation can be a significant addition to our understanding of monoaminergic neurotransmission. Monoamines, and especially serotonin, display several characteristics that separate them from other neurotransmitters. Their exocytosis is both spatially and temporally diffuse, and their effects can lead to long-term neuromodulation.^15,16^ A membrane-mediated pathway can facilitate these wide and diffuse effects, and can also provide a handle for potential pharmaceutical agents to modulate these effects.

Here we probe both the process of membrane attachment and the process of membrane fusion *in vitro*, as a function of the concentration of externally added serotonin. We probe the attachment, dissociation, and fusion of vesicles by Fluorescence Correlation Spectroscopy and TIRF microscopy. We note that the actual exocytosis process in live cells is complex. The proximity of the vesicles to the bilayer is regulated by the Rab family of proteins.^17^ The process is modelled by a series of calcium dependent maturation steps (docked **→** pre-primed **→** primed) before the actual fusion takes place where the pre-primed and primed states are reversible.^18,19^ While the intracellular process of exocytosis involve the cellular machinery of various proteins^20^, it is useful to ask if the membrane itself can play any role in this process. In our *in vitro* spontaneous fusion assay, we model this exocytosis process as reversible ‘docking’ and subsequent fusion of artificial lipid vesicles with supported lipid bilayers (SLBs). We use vesicles whose size^21,22^ and composition^23^ resemble those of the synaptic vesicles and probe if any of these steps are influenced by serotonin present in the solution. We note that the experimental conditions used in these experiments (e.g. externally added serotonin, absence of any regulatory proteins etc.) are different from those experienced by the serotonergic vesicles present in neurons. However, our study is performed in a pre-defined well-characterized lipid system, which can highlight any potential role that the lipid bilayer itself may play in real neurons. Additionally, we also probe the molecular mechanism underlying this effect by probing the serotonin-induced changes in the state of the lipid bilayers, using spectral imaging and molecular dynamics (MD) simulation tools.

Our results reveal a significant effect of monoamine neurotransmitters, especially serotonin, on the process of vesicular fusion. In real neurons, such a mechanism can provide the vesicular cargo with unique control on its own release.

## 2. Results

### 2.1 Serotonin and other monoamines promote vesicular association

We probe whether serotonin affects the association between small unilamellar vesicles (SUVs) of composition PC/PE/PS/Chl 3/5/2/5 (this molar ratio represents the composition of the major lipids present in the synaptic vesicles^23^) in 20 mM HEPES buffer (300 mM NaCl, pH 7.4) using Fluorescence Correlation Spectroscopy (FCS). FCS measures the diffusion of fluorescent particles, which would tend to slow down if the vesicles associate with each other. The vesicles were fluorescently labelled with Nile Red to make them amenable to FCS. The representative autocorrelation curves are shown in Fig. 1A. The timescale of decay of the FCS trace becomes longer in the presence of 5 mM serotonin indicating the formation of larger diffusing particles. The FCS traces were fitted with a three-dimensional diffusion model (eq 1) with two separate diffusing species corresponding to free and vesicle-bound Nile Red molecules respectively. We note that the vesicle sizes are likely to be heterogeneous, but a two-component model suffices for comparison. The residuals of the fits are shown in Fig. 1A. τ _D1_ corresponds to the diffusion of free Nile Red molecules, while τ _D2_ represents the vesicles. The average hydrodynamic radii (R_H_) of the vesicles (eq 4) at different concentrations of serotonin (Fig. 1B) is calculated from τ _D2_ using rhodamine dye (R_H_ = 0.58 nm) as a calibrant^24^. The R_H_ values obtained are 23.4 ± 2.5 nm, 36.9 ± 1.8 nm, 98.0 ± 6.9 nm, and 173.6 ± 50.6 nm in the presence of 0 mM, 0.1 mM, 0.5 mM, and 5 mM serotonin in the solution respectively. FCS measurements with Rhodamine-DOPE (Rh-PE) labelled (0.005 mol%) PC/PE/PS/Chl 3/5/2/5 vesicles were also carried out which showed a similar degree of association of the vesicles induced by 5 mM serotonin (Fig. S1). FCS measurements were repeated after dialyzing out the serotonin present in the solution in order to distinguish between reversible association and irreversible fusion. The hydrodynamic radius decreases nearly, but not completely, to the starting radius (Fig. 1B). This indicates that the size increase is mostly due to a reversible association of vesicles, with a minor fraction undergoing fusion.

**Fig. 1:**
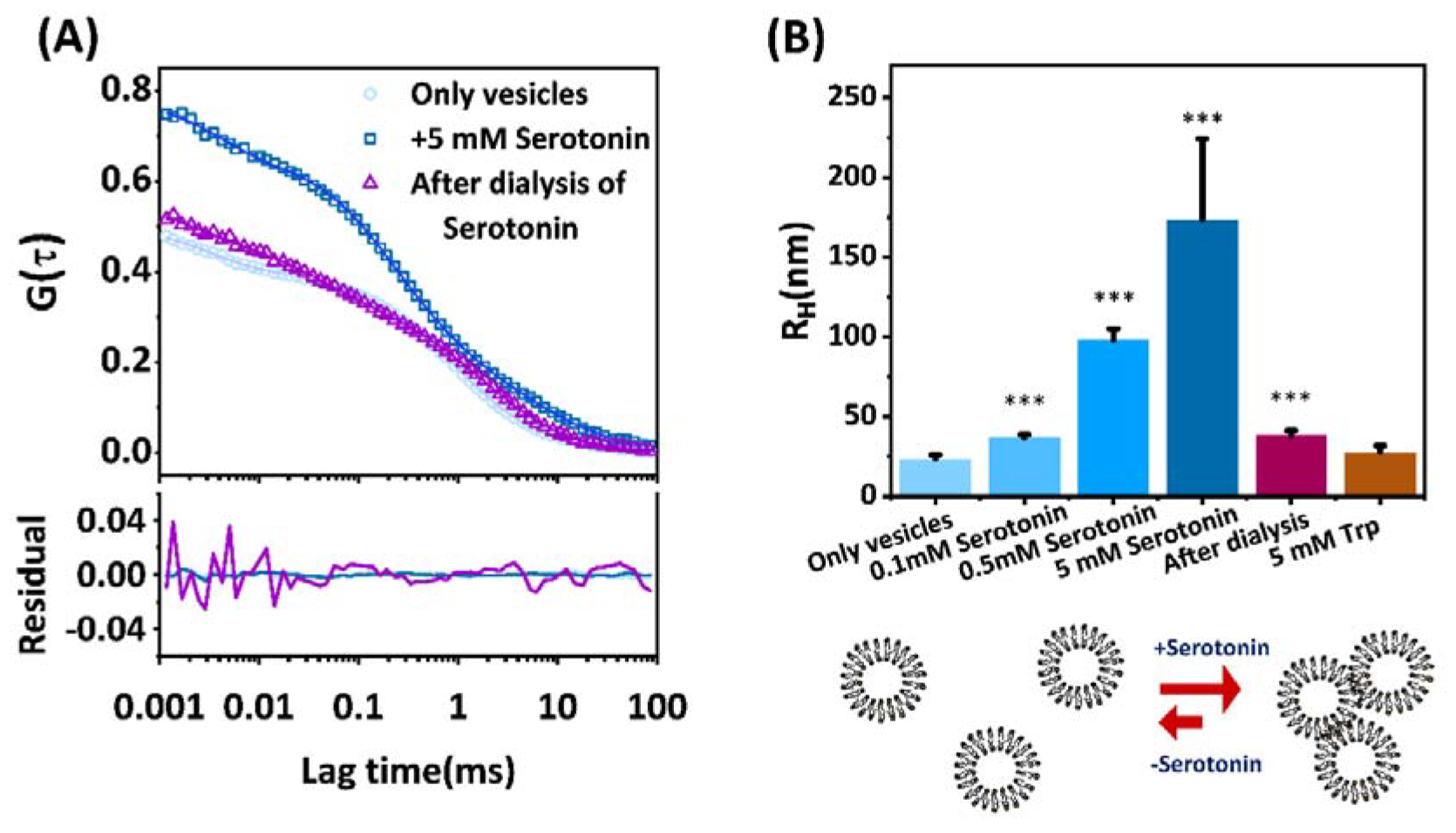
(A-B) Inter-vesicular association induced by serotonin. **(A)** Representative fluorescence autocorrelation curves of Nile Red labelled PC/PE/PS/Chl 3/5/2/5 vesicles (data: circles, squares, triangles) in 20 mM HEPES buffer (300 mM NaCl, pH 7.4) with their corresponding fits (solid lines) and residuals (solid lines at the bottom) in the absence of serotonin (sky blue), in the presence of serotonin (dark blue), and after dialysis (purple). **(B)** (upper panel) The average hydrodynamic radii of vesicles as a function of serotonin concentration and 5 mM Tryptophan (Trp); (lower panel) Cartoon representation of serotonin-induced reversible vesicular association. Mean and S.E.M are plotted. The distribution of the data was not normal (by Shapiro-Wilk test), therefore, a non-parametric Wilcoxon test with repeated measures correction was performed for significance measurements. ^*^p<0.05, ^**^p<0.01, ^***^p<0.001 compared to control (n= 6 or larger).

We also tested whether this effect is specific to serotonin, or whether other similar molecules can have similar effects. We performed FCS measurements in the presence of dopamine and norepinephrine (NE) (Fig. S2). The average hydrodynamic radius of the diffusing particles increases to 46.3 ± 3.3 nm in the presence of 5 mM norepinephrine and to 122.7 ± 10.2 nm in the presence of 5 mM dopamine (Fig. S2B). Therefore, dopamine and norepinephrine also have effects similar to serotonin, though the effects are somewhat weaker. The effect on vesicle association follows the order serotonin > dopamine > norepinephrine. We then tested the effect of tryptophan (Trp) which is structurally similar to serotonin (though it is not a monoamine) and is a precursor of serotonin^25,26^. The R_H_ does not change significantly in the presence of 5 mM Trp (Fig. 1B; no Trp: 23.4±2.5 nm, with 5 mM Trp: 27.5±4.2 nm; FCS traces are shown in Fig. S3).

Our results, therefore, show that serotonin and other monoamines specifically promote the association of lipid vesicles. However, the chemically related molecules, such as tryptophan, do not show such effects.

### 2.2 Enhanced association of vesicles to supported lipid bilayer promotes fusion

While FCS does not measure rate constants for binding, dissociation, and fusion of vesicles (k_ON_, k_OFF_, and k_FUS_ respectively), direct vesicular imaging using TIRF^27^ enables us to do so. It is important to measure the effect of serotonin on the association and fusion of vesicles with the SLB, which is relatively flat and is a better model for the plasma membrane. We note that the vesicular fusion process in living cells is more complex and involves a series of priming steps regulated by the SM family and SNARE proteins before the actual release takes place.^18,19^ For *in vitro* spontaneous fusion experiment used here, the vesicles that diffuse to the vicinity of the SLB can spontaneously bind to it, and then can either dissociate or irreversibly fuse with it (Fig. 2A). This process can be depicted by the following equation:

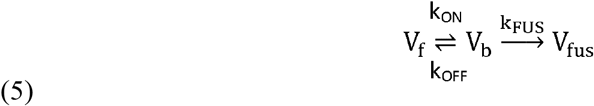

Here V represents vesicles, and the subscripts f, b, and fus denote the free, bound, and fused vesicles respectively. We considered only those events where the vesicles stayed on the SLB for at least 2 frames (50 ms). This effectively discards the vesicles which diffuse into the evanescent field but do not come in contact with the bilayer. We performed TIRF experiment (Fig. 2B) with Rh-PE labelled vesicles and unlabelled SLB of the same lipid composition (PC/PE/PS/Chl 3/5/2/5) at increasing concentrations of serotonin (0 mM, 0.1 mM, 0.5 mM, 2.5 mM, and 5 mM). The binding rate (R_ON_) does not change with serotonin (Fig. S5) as it is controlled by the collision frequency, which in turn is determined by the size and concentration of the vesicles. We note that the concentration of the vesicles is so low (500 fM) here that there is no significant binding between the vesicles in the aqueous phase before or during the experiment. We observe that the dissociation rate constant (k_OFF_) decreases by more than 10 times in the presence of 5 mM serotonin (Fig. 2C). The dissociation rate constant of vesicles without serotonin is 5.50 ± 0.13 s^1^ and it changes to 4.48 ± 0.94 s^-1^, 2.45 ± 0.77 s^-1^, 0.55 ± 0.15 s^-1^, 0.46 ± 0.09 s^-1^ in presence of 0.1 mM, 0.5 mM, 2.5 mM, and 5 mM serotonin respectively.

**Fig. 2:**
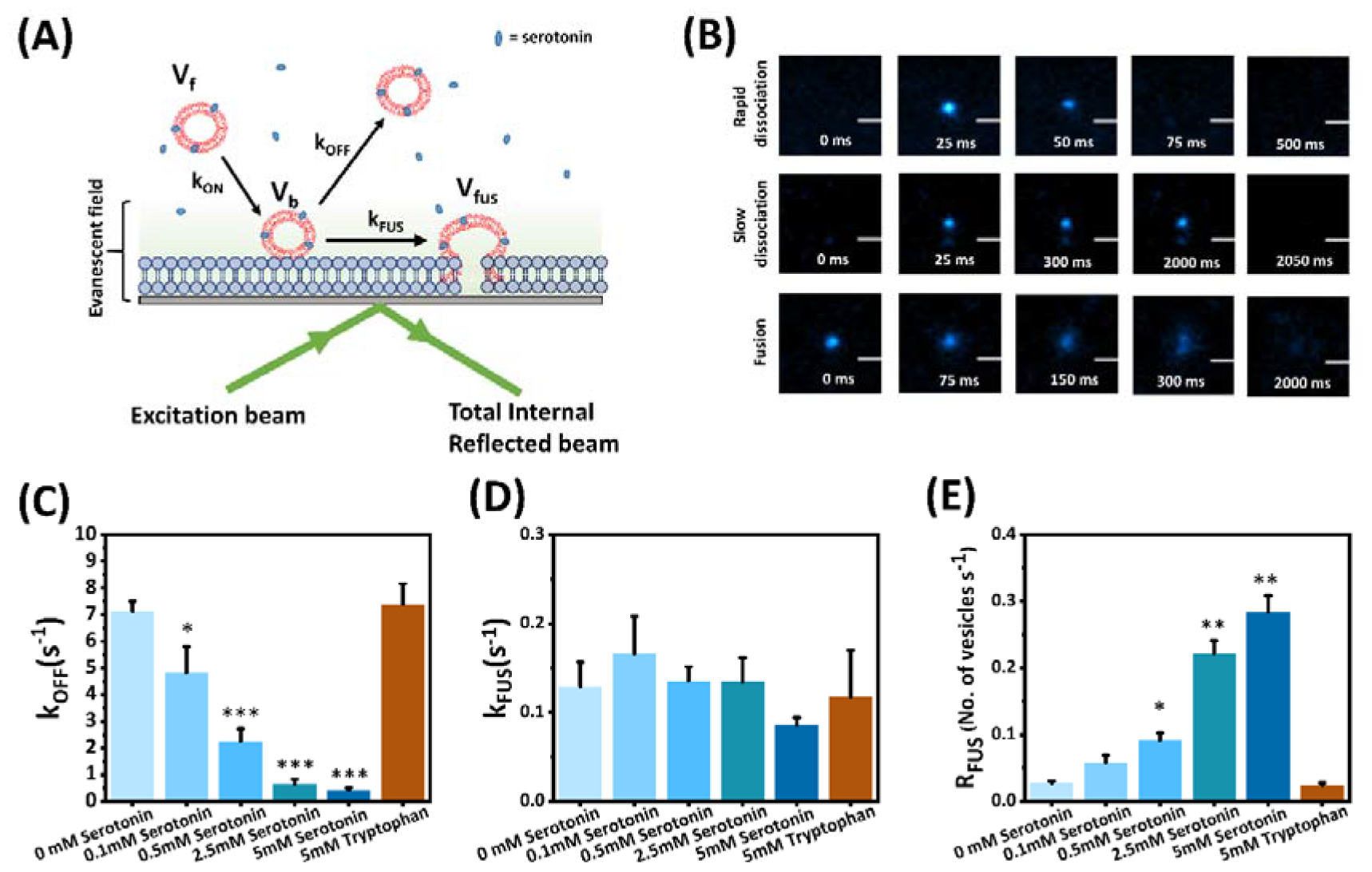
Serotonin-induced association and fusion of vesicles to SLBs. **(A)** Schematic representation of vesicle adhesion and fusion to Supported Lipid Bilayers (SLB) probed using a Total Internal Reflection Fluorescence (TIRF) microscope (not to scale). The vesicles in the solution come in contact with the bilayer with a rate constant k_ON_. The bound vesicles can revert back to the solution with a rate constant k_OFF_ or fuse with the bilayer with a rate constant k _FUS_. The vesicles are fluorescently labelled with 0.08 mol% Rh-PE. **(B)** Different types of events observed in TIRF imaging: (upper panel) rapid binding and dissociation of vesicle on bilayer, (middle panel) vesicle adhered on the bilayer for a longer time, (lower panel) fusion of the vesicle with the bilayer. **(C), (D)** and **(E)** k _OFF_, k_FUS_, and Rate of fusion (R_FUS_) as a function of serotonin concentration (0 mM, 0.1 mM, 0.5 mM, 2.5 mM, 5 mM) and 5 mM tryptophan. Mean and S.E.M are plotted. This dataset passed tests for normality (Shapiro-Wilk test and Kolmogorov-Smirnov test, p>0.05) and equality of variances (F-test for variances, p>0.05). Therefore, ANOVA tests with repeated measures correction were performed to test the significance. ^*^p<0.05, ^**^p<0.01, ^***^p<0.001 compared to control (only vesicles) (n=5).

The rate constant for the fusion step (k_FUS_), which represents the probability of fusion per unit residence time, does not change significantly (Fig. 2D). However, the total rate of fusion (R_FUS_) is the product of k_FUS_ and V_b_ (see eq.13 in SI section 1.6), and V_b_ increases as k_OFF_ decreses. So while k_FUS_ remains unchanged, k_OFF_ becomes much smaller, resulting in an increase in the net rate of fusion (R_FUS_, Fig. 2E). R_FUS_ increases from 0.028 ± 0.008 vesicles s^-1^ to 0.049 ± 0.027 vesicles s^-1^, 0.085 ± 0.033 vesicles s^-1^, 0.237±0.053 vesicles s^-1^ and 0.272 ± 0.071 vesicles s^-1^ in presence of 0.1 mM, 0.5 mM, 2.5 mM, and 5 mM respectively. We observe that tryptophan does not have any substantial effect on the dissociation rate constant (k_OFF_), k_FUS_, or rate of fusion (R_FUS_) of the vesicles (Fig. 2C, Fig. 2D, and Fig. 2E respectively).

### 2.3 Serotonin-induced vesicle association is not driven by electrostatics

We then probed the possible molecular mechanisms driving this enhanced association between vesicles. Since serotonin is a positively charged molecule at pH 7.4 (Fig. S6) and the membrane is negatively charged (because of the presence of POPS), it is plausible that the vesicular association behavior induced by serotonin lowers the electrostatic repulsion between the bilayers and drives the association. Such electrostatic repulsion would be screened at higher ionic strength solvents, so we probed the effect of 5 mM serotonin at different concentrations of NaCl (0 mM, 150 mM, and 300 mM NaCl) (Fig. S7A). We observe an increase in the R_H_ of vesicles at all concentrations of NaCl (0 mM NaCl: 21.4 ± 1.2 nm to 110.8 ± 21.1 nm, 150 mM NaCl:17.9 ± 0.4 nm to 77.6 ± 6.6 nm, 300 mM NaCl: 23.4 ± 2.5 nm to 173.6 ± 50.6 nm). The increase is not correlated with the salt concentration. Furthermore, if electrostatic effects are dominant, they should not enhance the association between neutral vesicles. We observe that for POPG (100% negatively charged headgroups), PPC111 (33% negatively charged headgroup), and POPC (neutral headgroup) vesicles, 5 mM serotonin induces qualitatively similar increases in the R_H_ (Fig. S7B, POPG: 26.1 ±2.9 nm to 73.6±5.0 nm, PPC111: 19.2±2.1 nm to 46.4±2.1 nm, POPC: 19.5 ± 0.2 nm to 76.4 ± 4.9 nm). We also probed the effect of 5 mM dopamine and tryptophan on POPC (neutral headgroup) (Fig. S4A) and POPS (negatively charged headgroup) (Fig. S4B) lipid vesicles. In both cases, dopamine shows significant vesicle association but tryptophan does not. All these observations suggest that electrostatics does not play a major role in driving vesicular association.

### 2.4 Serotonin increases the average area occupied by a lipid molecule

An alternative hypothesis is that serotonin promotes association by creating defects/disorder in the membrane which in turn leads to attractive hydrophobic interactions between vesicles. Using ^2^H-NMR spectroscopy, we have observed previously^8,10^ that serotonin increases disorder in the lipid chain. ^2^H-NMR experiments with multilamellar vesicles (of composition PC-d31/PE/PS/Chl 3/5/2/5) provided the order parameter (S_CD_) for each carbon along the lipid chain (reproduced with permission from Gupta et al. ^10^ in Fig. S8A). In the presence of 10 mol% serotonin, the order parameter profile homogeneously decreased along the entire carbon chain. The decrease in the lipid chain order decreases the average chain length (<L_c_>) of the lipid from 16.47Å to 16.21 Å. Assuming that the lipid density remains unaltered, we calculate the average lateral area occupied per lipid molecule (<A>, see eq. 14) of POPC in PC/PE/PS/Chl 3/5/2/5 vesicles in the presence of 0 and 10 mol% serotonin respectively (Fig. S8B), and observe that <A> increases from 51.5 Å^2^ to 52.3 Å^2^ in presence of serotonin.

The increase in the lipid chain disorder and the lateral area occupied by lipid molecules suggests the formation of local defects in the bilayer. Since the bilayer is in equilibrium with water near its headgroup region, this can increase the unfavorable penetration of water in the lipid-water interface of the bilayer. Association of two lipid bilayer can reduce the water concentration at the inter-bilayer interface and therefore also the energetically unfavorable penetration of water into the bilayer.

### 2.5 Serotonin promotes water penetration into the bilayer

To investigate water penetration into the bilayer at a molecular level, we performed all-atom molecular dynamics (MD) simulations of both the neutral POPC membrane and negatively charged POPS membrane in the absence and the presence of 2 and 20 serotonin molecules respectively, with 200 lipid molecules. Considering the partition coefficient of serotonin in lipids this is in the range of membrane concentrations attained by adding ∼0.5 to ∼5 mM serotonin in the solution, as estimated previously by us^3,8^. We calculated the z-axis distance between the lipid bilayer center and water molecules using 5000 snapshots from the second halves of the 1000 ns simulations. The number of water molecules along the POPC and POPS bilayer normal is plotted in Fig. 3A and Fig. 3B respectively. We notice a significant increase of water molecules inside the core of the hydrophobic part of the lipid bilayer in the presence of serotonin molecules. The maximum difference is observed around 0.5-1 nm from the centre of the bilayer. We note that POPC and POPS are simple model lipid membrane system which does not mimic the vesicular lipid composition (PC/PE/PS/Chl 3/5/2/5). However, we verified that serotonin also promotes vesicular association for POPC and POPS lipid vesicles (Fig. S7B). We, therefore, consider this phenomenon to be general and independent of the lipid composition. We note that the magnitude of water penetration correlates with the magnitude of vesicular association in POPC and POPS systems (Fig. S7B and Fig. 3A, B).

**Fig. 3:**
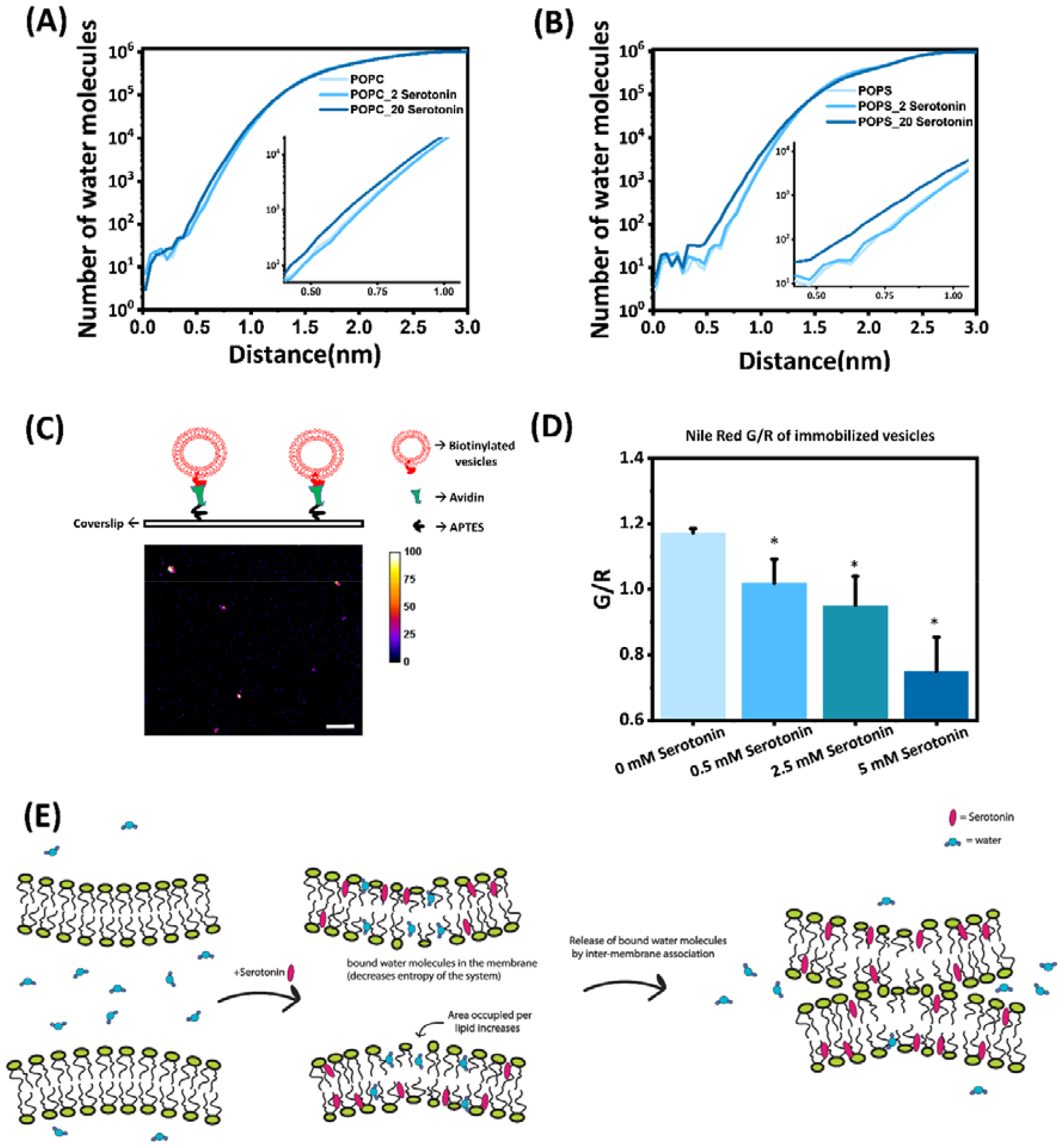
Effect of serotonin on the penetration of water molecules in a bilayer. **(A) and (B)** MD simulation of the number of water molecules as a function of the distance from the centre of the **(A)**POPC and **(B)**POPS bilayer in the absence and presence of 2 and 20 serotonin molecules. The inset shows a magnified region at distances where serotonin has maximum effect. **(C)** (upper panel) Cartoon representation of the method of immobilization of vesicles on glass coverslips. (lower panel) A representative confocal fluorescence image of Nile Red labeled PC/PE/PS/Chl 3/5/2/5 vesicles immobilized on the glass surface. **(D)** Nile Red peak shift (Green/Red or G/R ratio) of vesicles obtained from spectral imaging of vesicles as a function of the concentration of serotonin (0 mM, 0.5 mM, 2.5 mM, 5 mM). Mean and S.E.M are plotted. The dataset passed tests for normality (Shapiro-Wilk test and Kolmogorov-Smirnov test, p>0.05) and equality of variances (F-test for variances, p>0.05. Therefore, repeated measures t-tests were performed to check the significance level. ^*^p<0.05, ^**^p<0.01, ^***^p<0.001 compared to control (only vesicles) (n=4). **(E)** Cartoon representation of the mechanism of serotonin-induced inter vesicular membrane association. The vesicle association is mainly driven by hydrophobic attraction induced by serotonin due to an increase in local membrane defects leading to increased exposure of the lipid chain to the aqueous phase.

We also test the water penetration of the bilayer experimentally. We incubated PC/PE/PS/Chl 3/5/2/5 vesicles with Nile Red, whose fluorescence emission spectrum is sensitive to water ingress into the membrane^28,29^. We quantified the water ingression in terms of the G/R ratio of Nile Red, where ‘G’ and ‘R’ denote the total emission intensity in the wavelength range 570-600 nm and 610-640 nm respectively. It is important to examine the water content of the bilayer without allowing the vesicles to associate, as the association may lower the water content by drying the inter-bilayer interface. We immobilized the vesicles on glass coverslips (Fig. 3C) using biotin-streptavidin chemistry to prevent association. The fluorescence spectrum of Nile Red-labeled immobilized vesicles was recorded by spectral imaging with a confocal fluorescence microscope. We observed a dose-dependent decrease in the G/R ratio with an increase in the concentration of serotonin (Fig. 3D). The G/R ratio of vesicles in HEPES buffer (pH 7.4) decreases from 1.16 ± 0.06 to 1.08 ± 0.08, 0.95 ± 0.05, and 0.83 ± 0.06 in the presence of 0.5 mM, 2.5 mM, and 5 mM serotonin respectively. This shows that the water ingress into the vesicular membrane increases with an increase in the concentration of serotonin. The Nile Red spectra remained unaltered by 5 mM tryptophan (Fig. S9A-B). We note that a similar increase in water penetration by another monoamine, dopamine, has been previously shown experimentally using X-ray diffuse scattering (XDS).^6^

## 3. Discussion

Two major steps in the process of regulated exocytosis of synaptic vesicles that can involve active participation of the membrane are (1) attachment of vesicles to the plasma membrane, and (2) fusion of the attached vesicles with the plasma membrane.^30^ While both of these steps in a cell are thought to be primarily driven by proteins (Rab and SNARE family proteins respectively^20,31^), any change in membrane properties that facilitate either of these steps can have a significant effect on exocytosis. In this work, we have quantified the attachment and fusion steps separately in an artificial lipid vesicle system and probed the effect of serotonin on each of these steps. From FCS measurements, we found that serotonin increased association between the vesicles (Fig. 1A and 1B). TIRF measurements show that the serotonin-induced increased duration of association enhances its probability of fusion with the supported lipid bilayer (Fig. 2). The observed increase of an order of magnitude in the rate of fusion of vesicles is driven majorly by the increased duration of association between the vesicle and the supported lipid bilayer, and not by a lowering of the free energy barrier for fusion. We note that other monoamine neurotransmitters (dopamine, and norepinephrine) also induce vesicular association (Fig. S2). Therefore, these processes can be general for monoamines.

Several repulsive and attractive forces can govern the association of vesicular membranes.^32^ We find that the vesicular association is not driven by electrostatics. (Fig. S7). The other possibility can be an enhanced hydrophobic interaction^32^ induced by serotonin. In the case of free bilayers, the hydrophobic tails of the lipid molecules are shielded by the hydrophilic head groups which mask the hydrophobic interaction between two bilayers. When bilayers are subjected to lateral stress, the surface area per lipid increases, which leads to the exposure of the hydrocarbon tail to the aqueous phase. An association between two bilayers can reduce this unfavorable exposure. Serotonin decreases the lipid chain order of lipid bilayers suggesting an increase in surface area^8,10^ (Fig. S8). Also, it has been shown to alter the phase equilibrium of a ternary mixture of lipids, driving the system to a more disordered state.^7^ Therefore, serotonin can increase the water penetration of the bilayer. The increased penetration of the hydrophobic region of the bilayer by water has an entropic cost. Vesicle association, in turn, can expel the water from the inter-bilayer region. Less water near the interface would in turn cause a drying of the hydrophobic region of each of the bilayers. This would lower the free energy of the bilayers, stabilizing the association between the bilayers. Such effects are well known in other contexts. For example, Ca^2+^ drives vesicular association by causing a local decrease in lipid packing.^32,33^ MD simulations (Fig. 3A-B) and Nile Red spectral imaging (Fig. 3C-D) showed increased penetration of water in the bilayer. We infer that this increased hydrophobic interaction can have a significant contribution to drive vesicular association, as elaborated in Fig. 3E. We note that dopamine has also been shown to increase the penetration depth of water molecules in the lipid bilayer^6^ which corroborates with its effect on vesicular association observed by us (Fig. S2). Though our results demonstrate a possible mechanism of serotonin-induced bilayer association, our data cannot exclude other serotonin-induced alterations in the repulsive and attractive forces (e.g. repulsive thermal fluctuation force, headgroup overlap force, protrusion force and attractive Van der Waals forces and depletion forces) which too can contribute to vesicular association.

## Conclusion

The mechanical properties of the lipid bilayer impact a large fraction of all biochemical functions of the living cell. It controls major processes such as endo- and exocytosis, membrane protein conformation and dynamics, response to mechanical stimuli, motility, intracellular transport, and neuronal plasticity. Any agent that can change its properties has the potential to affect a multitude of cellular functions. Our results show that monoamines can act as such agents. The relationship between monoamine concentration and the propensity of association and fusion of artificial lipid vesicles reported here can potentially have major biological significance, such as for exocytosis. Exocytosis is the critical initial step in all chemical neurotransmission in the brain and its modulation by the neurotransmitter brings in an additional control pathway to the chemical network of neurotransmission. The current understanding of vesicular exocytosis assumes that the neurotransmitter itself has no control over exocytosis. Our results can potentially change that view. Also, transient serotonin concentrations during and immediately after exocytosis, both in the synaptic cleft and in the plasma membrane at the presynaptic active zone locations, are expected to be high. This can facilitate binding and exocytosis of other vesicles in that region. Also, processes other than exocytosis, such as ultrafast endocytosis that is coupled to exocytosis^34^, can be facilitated by a transient change of the mechanical properties of the membrane at locations of recent exocytosis.

In addition to neurotransmission in the brain, monoamines are known to modulate a large range of cellular functions. Serotonin is present at high concentrations in many organs, such as the enteric system, the endocrine system, and in the developing embryo. So a direct perturbation of the mechanical properties of the lipid bilayer by monoamines points to a set of chemical pathways by which an organism may control a multitude of functions. While specific receptors provide a tightly regulated response to the alteration of monoamine concentrations at specific locations, the membrane-mediated pathway can provide a more distributed and longer-lasting system-wide response to changes in monoamine levels. A failure to take into account this additional pathway may have limited our understanding of the effects of monoamines in different organs of our body, at different stages of development.

## Abbreviations

(AFM): Atomic Force Microscopy
(SLB): Supported lipid bilayer
(NMR): Nuclear Magnetic Resonance
(TIRF): Total Internal Reflection Fluorescence
(Trp): Tryptophan
(SUVs): Small unilamellar vesicles
(MD): Molecular dynamics

## Data Availability

Detailed experimental procedures having Materials and Methods, and all supplementary figures (Fig. S1-S11) are provided in ESI.

## Author contributions

DSR performed most of the experiments and analysed the data. AG and VV helped perform the FCS measurements. PK and MSL performed the simulations. DSR, PK, MSL and SM co-wrote the manuscript. SM conceptualized the project.

## Acknowledgments

This work was supported by the Department of Atomic Energy, Government of India, with a grant to SM provided under project no. RTI4003. We thank Prof. Daniel Huster and Prof. Madan Rao for fruitful discussions regarding this work.

