## Supporting information for "Serotonin Promotes Vesicular Association and Fusion by Modifying Lipid Bilayers"

*Corresponding author

**1. Materials and methods**

**1.1. Materials:**

The lipids POPC (16:0-18:1 PC), POPE (16:0-18:1 PE), POPS (16:0-18:1 PS), POPG (16:0-18:1 PE), 1,2-dioleoyl-sn-glycerol-3-phosphoethanolamine-N (lissamine rhodamine B sulfonyl) (18:1 Liss Rh-PE) were purchased from Avanti Polar Lipids Inc. (Alabaster, USA). Cholesterol and Na-HEPES were purchased from Sigma Aldrich (St. Louis, MO). Serotonin-hydrochloride was purchased from Merck (Darmstadt, Germany). Sodium chloride and Potassium Chloride were purchased from Merck Life Science Private Limited (India). Chloroform AR graded was purchased from S.D. Fine Chemical Ltd. (India). A lipid extruding kit and Nucleopore Tracketched polycarbonate membranes of 50 nm pore diameter were bought from Avanti Polar Lipids (Alabaster, AL). Biotech CE Tubing:100 kDa MWCO dialysis membrane was purchased from Spectrum Laboratories, Inc. (MA, USA). All the chemicals were used without any additional purification. Flat supported lipid bilayers were formed on glass coverslips of refractive index-1.78 (Olympus APO100X0HR). All the experiments were performed using milli-Q water deionized to 18.2 MΩ cm^-1^ obtained from a milli-Q gradient system (Millipore, Germany).

**1.2. Preparation of Small Unilamellar Vesicles (SUVs):**

Small Unilamellar Vesicles (SUVs) were prepared following a previously published protocol^1^. Briefly, the lipids POPC, POPS, POPG, POPC/POPE/POPS/Cholesterol in the molar ratio of 3/5/2/5 (PC/PE/PS/Chl 3/5/2/5), and POPC/POPG/cholesterol in the molar ratio of 1/1/1 (PPC111) were prepared by weighing the powders in the respective molar ratio and dissolving them in Chloroform solvent. The solvent was then evaporated by purging iolar Argon gas while rotating the vial to prepare a thin film of lipids. The lipid film was kept in a vacuum desiccator for ~24 hours to completely remove organic solvents and confirm the film is dry. The dried film was then rehydrated using HEPES buffer (20 mM HEPES, pH 7.4, 0/150/300 mM NaCl) to make a 2.5 mg/ml lipid suspension. The solution was vortexed properly for ~20 minutes to form multilamellar vesicles (MLVs) of the corresponding lipids. The MLVs were passed through an extruder (Avanti Polar Lipids Inc., Alabaster) containing a polycarbonate membrane of 50 nm pore size (covered by two 10 mm filter supports on each side of the membrane) using 1000 µL vacuum-sealed syringes (Hamilton, Avanti Polar lipids Inc.) The extrusion was carried out at 50 °C. This led to a clear solution containing uniform SUVs.

**1.3. Preparation of Supported Lipid Bilayers (SLBs):**

The SLBs were formed on high-refractive glass coverslips using the vesicle fusion method^2^. 15 µl of the SUVs having bilayer composition PC/PE/PS/Chl 3/5/2/5 were added with 55 µL of HEPES buffer (all were preheated to 65°C in a water bath) in the middle of a chamber, glued to the freshly cleaned (both piranha solution and plasma-treated) and dried glass coverslip with 10 µl of 100 mM CaCl_2_ solution and incubated for 1 h at 65°C in a water bath and slowly cooled down to room temperature. The vesicles fuse to form a bilayer in this condition. The lipid bilayer was then rinsed extensively with HEPES buffer to remove the unfused vesicles. The bilayer was then kept for equilibration for 30 min before starting the experiment.

**1.4. Fluorescence Correlation Spectroscopy (FCS):**

We used FCS for measuring the size (hydrodynamic radius, R_H_) of the vesicles^3,4^. 10 nM Nile Red dye was incubated for 20 minutes with the vesicle solution having ~10 nM concentration of vesicles. The vesicle solution was then deposited on a glass coverslip and FCS measurements were performed. Each FCS measurement was repeated more than or equal to 6 times. The results shown are the average ± SEM of the 6 or more replicates. These measurements were performed using a home-built FCS instrument^4^. Briefly, a 488 nm laser beam was expanded and collimated using a 1:4 telescope set-up. The collimated excitation beam is then focused into the sample using an apochromatic 60X water immersion objective with a numerical aperture of 1.2 (Olympus, Center Valley, PA). The fluorescence was collected using the same objective and separated from the excitation beam using a 500 nm LP dichroic mirror (Chroma Technology, Rockingham, VT). The emission beam was then focused onto a 15-μm core-diameter optical fibre after filtering through a 607/70 nm band pass emission filter (Chroma Technology, Rockingham, VT). The fibre was used as a confocal pinhole to reject the out-of-focus fluorescence. The fluorescence was detected by a single-photon avalanche photodiode (PerkinElmer, Waltham, MA) which was connected to the other end of the fibre. The data were collected and processed using a hardware correlator (PicoHarp 300; PicoQuant, Berlin, Germany). A two-component, three-dimensional diffusion model with a triplet component was used (equation 1) to fit the FCS data using Origin 6.0 software.

G(τ) = $\left( \begin{aligned} \left( 1-f+f*e^{-\tau/{\tau_{t}}} \right) \\ \bar{(1-f}) \end{aligned} \right)*[\frac{g1}{(\left( 1+\frac{\tau}{\tau_{D1}} \right)*\sqrt{\left( 1+a^{2}*\frac{\tau}{\tau_{D1}} \right)})}+\frac{g2}{(\left( 1+\frac{\tau}{\tau_{D2}} \right)*\sqrt{\left( 1+a^{2}*\frac{\tau}{\tau_{D2}} \right)})}]+bl.$	(1)

Where, $a=\frac{w_{0}}{z_{0}}$

Here, G(τ) is the autocorrelation function at lag time τ, f is the fraction of the triplet component and τ_t_ is the corresponding triplet lifetime. Also, g1 and g2 are the amplitudes of the correlation function corresponding to the two diffusing components (the free Nile red and Nile red bound to vesicles), τ_D1_ and τ_D2_ are the corresponding diffusion times, a is the structure parameter for the optical probe volume (assumed to be a Gaussian ellipsoid), and bl denotes the background signal. z_0_ and w_0_ are the length and width of the focal volume respectively. Free Rhodamine B dye was used as a standard to calibrate the instrument. We obtained a τ_D_ of 29 μs for free Rhodamine B in the HEPES buffer (pH 7.4). The R_H_ for free Rhodamine B was considered to be 0.58 nm^5^. The diffusion times of the Nile red-labelled vesicles(τ_D2_) were converted to R_H_ by comparing their diffusion times with that of free Rhodamine B in solution according to equation 4.

$\tau_{D}=\frac{w_{0}^{2}}{4D}$	(2)

$D=\frac{K_{B}T}{6\piɳR_{H}}$(Stoke-Einstein’s equation)	(3)

Therefore, $R_{H(vesicle)}=\frac{\tau_{D(vesicle)}}{\tau_{D(Rhodamine B)}}*R_{H(Rhodamine B)}$	(4)

**1.5. Total Internal Reflection Fluorescence (TIRF) imaging of single vesicle association and fusion:**

We used a home-built TIRF microscope^6^ for imaging single vesicle association and fusion to the SLBs. The samples were excited with a 543 nm He–Ne continuous laser (Melles Griot, 25-LGR-393-230). The beam was expanded from 2 mm to 20 mm using a 1:10 Gaussian telescope set-up. The collimated beam (power ∼1.3 mW) was focussed at the back aperture of the objective lens by using a 50 cm lens kept on a translation stage. The laser was guided into the objective from the backport of the microscope (Ti-Eclipse, Nikon, Japan). A 565 nm dichroic mirror was used to separate the fluorescence from the excitation laser. Fluorescence was collected using a band-pass emission filter (BA 577–633 nm, Nikon, Japan) and focused on an electron-multiplying CCD camera (ANDOR iXON, DV887ECS-UVB) mounted on the side port of the microscope. SLBs of composition PC/PE/PS/Chl 3/5/2/5 were prepared on a high-refractive glass coverslip as described in section 1.3 and kept on top of the microscope. The SUVs having lipid composition PC/PE/PS/Chl 3/5/2/5 with a 0.08-mole fraction of Rh-PE were prepared as described earlier. The vesicle solution was diluted to a concentration of 500 fM and 140 µl of the solution was added to the chamber containing SLB. The single fluorescent vesicle dynamics on the bilayer are monitored by acquiring TIRF images continuously with a frame rate of 40 Hz. Each measurement was repeated 5 times. The results shown are the average ± SEM of the 5 replicates.

**1.6. Analysis of vesicular dynamics on SLB from TIRF imaging experiment:**

The adhesion and fusion of vesicles on SLB can be depicted by equation (5) considering the observable for our measurement to be the fluorescent vesicles.

k_ON_

$V_{f}\rightleftharpoons V_{b}\underset{\to}{k_{\mathrm{FUS}}}V_{\mathrm{fus}}$	(5)

k_OFF_

V_f_ is the state of vesicles when they are freely moving in solution. V_b_ is the state where the vesicles are physically bound to the bilayer. The bound vesicles can dissociate to the V_f_ state or can irreversibly convert to the V_fus_ state where the vesicles fuse with the bilayer. By TIRF imaging, we can distinguish between fusion and dissociation events. The dissociation of vesicles will give rise to a single-step disappearance whereas fusion will lead to a gradual spreading of the fluorescence due to diffusion of the Rh-PE lipids in the bilayer concomitant with the disappearance of the vesicles (Fig 2B). The transition from V_b_ to V_fus_ state is a non-equilibrium process. However, the number of vesicles undergoing fusion is very low compared to the total number of vesicles in the solution. Given the concentration and volume of the vesicle solution we used, only 2% of the total vesicles can collide with the SLB calculated using the Ward-Tordai equation^7^ considering Fick’s law of diffusion for the vesicles in liquid. Therefore, we assume V_f_ and V_b_ to be time-invariant.

Therefore,

Rate of binding = R_ON_ = k_ON_[V_f_] = Number of vesicles that appear near the SLB per second	(6)

Rate of fusion (R_FUS_) = k_FUS_ [V_b_] = Number of vesicles fused per second	(7)

Therefore, k_FUS_ = $\frac{Number of vesicles fused per second}{The average number of vesicles bound to SLB}$	(8)

Again, Rate of dissociation = R_OFF_ = k_OFF_[V_b_]	(9)

This parameter is not observable in the TIRF imaging experiment

So, we define

R_2_=k_2_[V_b_]=(k_OFF_+k_FUS_)[V_b_]	(10)

Where	k_2_ = $\frac{1}{The average residence time of vesicles on the SLB}$	(11)

Using equations (8), (10) and (11), we get

k_OFF_ = k_2_-k_FUS_	(12)

Again, the overall rate of the process (R_FUS_) = Rate of formation of V_fus_

= k_FUS_[V_b_]

= k_FUS_*$\frac{k_{\mathrm{ON}}}{k_{\mathrm{OFF}}}$*[V_f_]	(13)

**1.7. Correction of error in the calculation of k_ON_ and k_FUS_**

The time resolution of our instrument is about 25 ms. So, we possibly undercount the number of vesicles that appear on the SLB which gives an error in the calculation of k_ON_ and k_FUS_. To correct this, we calculate the probability of non-detectable events (defined as ‘r’) for a particular average residence time(t_av_) of vesicles considering the distribution of residence time to be exponential (Fig S17) as the dissociation from the flat bilayer is expected to follow a 1^st^ order rate kinetics.

So, r=$\frac{\int_{0}^{25} e^{\frac{-t}{t_{av}}}dt}{\int_{0}^{\infty} e^{\frac{-t}{t_{av}}}dt}$	(14)

Therefore, (k_ON_)_corrected_ =	$\frac{{(k_{\mathrm{ON}})}_{\exp}}{(1-r)}$	(15)

And, (k_FUS_)_corrected_ = (k_FUS_)_exp_*(1-r)	(16)

The experimentally observed k_ON_ and k_FUS_ are plotted in Fig S16A and Fig S16B respectively. The k_ON_ and k_FUS_ are corrected using eq 15 and eq 16 respectively and plotted in Fig S16C and Fig S16D respectively.

**1.8. Calculation of area occupied per lipid molecule from ^2^H NMR spectra of deuterium-labelled lipid vesicles**

Assuming a cylindrical model for lipids, the average lateral area occupied by lipid molecules(<A>) can be calculated by dividing the lipid chain volume by the lipid chain length (<L_c_>). Every methylene group takes up a volume of 26.5 Å^3^, for two chains it will be 53 Å^3^ per carbon.^8,9^ Every terminal methyl group takes up ~53 Å^3^.^9^ The first carbons are not included in the calculations.^9^

Therefore, <A>= ((n x 53Å^3^) + (2 x 53Å^3^))/ <L_c_> Å = 63.1 Å^2^	(14)

Where, n is the number of methylene groups in the lipid carbon chain and <Lc> is obtained from the order parameter values obtained from the ^2^H NMR spectra^10^).

**1.9. Measurement of Steady-state fluorescence spectra of Nile red**

Steady-state spectra of Nile red in PC/PE/PS/Chl 3/5/2/5 lipid vesicles with and without tryptophan were recorded in Fluorolog 3 (Horiba, Jobin Yvon Technology) by exciting at 532 nm. The emission was collected in the 560–700 nm spectral range. The fluorescence spectrum of Nile red is sensitive to the polarity of the membrane and shifts toward the red edge in a more polar environment.^11,12^ We quantified the water ingress in the membrane in terms of the G/R ratio where ‘G’ and ‘R’ denotes the total emission intensity in the wavelength range 570-600 nm and 610-640 nm respectively.

**1.10. Immobilization of vesicles on glass coverslips**

The vesicles were immobilized on glass coverslips using a well-established protocol^13^ with some modifications. Briefly, Corning glass coverslips were treated with Piranha solution for 1 hour. The coverslips were rinsed with water and ethanol and subsequently cleaned with Oxygen plasma for 30 minutes. This creates functional hydroxyl groups on the surface of the glass coverslips. The coverslips were then treated with 2% v/v APTES in ethanol for 4 hours to covalently link the APTES on the glass surface. The coverslips were rinsed with ethanol and then with buffer. After that, 0.4 mg/ml Avidin solution (dissolved in HEPES buffer, pH 7.4) was added and kept at 4°C overnight. In this step, Avidin binds with the free NH_2_ group of APTES. Unbound Avidin molecules were washed off by the HEPES buffer. 10 nM vesicles having 2% (w/w) Biotin-PE lipids were added and incubated for 4 hours to attach the vesicles with the avidin-coated coverslips through Biotion-Avidin interaction. The unbound vesicles were washed out using HEPES buffer.

**1.11. Confocal fluorescence imaging of immobilized vesicles on glass coverslips**

The Nile red spectra from immobilized vesicles were obtained by performing confocal imaging in the lambda stacking mode on the commercial LSM 880 (Carl Zeiss, Jena, Germany). The immobilized vesicles were incubated with 100 nM Nile Red in HEPES buffer (pH 7.4) for 20 minutes. The unbound Nile red and Nile red attached to the glass surface were washed off by the HEPES buffer multiple times. The spectral imaging of Nile Red was performed using a 543 nm excitation light from an Argon laser source focused onto the sample by a Zeiss C-Apochromat 40x, NA 1.2, water immersion objective. The fluorescence emission was collected by the same objective and separated from the excitation light by an MBS 543 dichroic mirror and sent to a GaAsP detector through a monochromator with a resolution of 10 nm. Lambda stacks were independently acquired from 560 to 670 nm at an interval of 10 nm with the help of ZEN imaging software.

**1.12. Measurement of water content in a bilayer using MD simulation**

To measure the number of water molecules penetrating lipid bilayers, 1000 ns all-atom MD simulations in NPT ensemble in Amber22 software were performed, starting from the equilibrated lipid bilayer models with and without serotonin obtained in our previous study^10^. Lipid14 force field^14^ was used to describe lipids, which was combined with a four-point OPC water model^15^ and Na+ and Cl- ions^16^, optimised for this water model, to neutralise the charge of the system and provide an ionic concentration of about 150mM. Obtained periodic boundary conditions boxes were equal to approximately 8.01x8.01x10.66nm (14458 water molecules) and 7.58x7.58x10.72nm (12574 water molecules) for 200 POPC and POPS lipid bilayers, respectively. Three variants of lipid compositions were studied: (i) without any serotonin molecules and with (ii) 2, and (iii) 20 serotonin molecules, resulting in total concentrations of approximately 0, 4.86mM, and 48.6mM for POPC, and 0, 5.39mM, and 53.9mM for POPS bilayers respectively. It should be noted that for the majority of the simulation time, serotonin molecules are bound or embedded in the lipid bilayer, therefore, the effective concentration is significantly lower.

**Supplementary Figures**

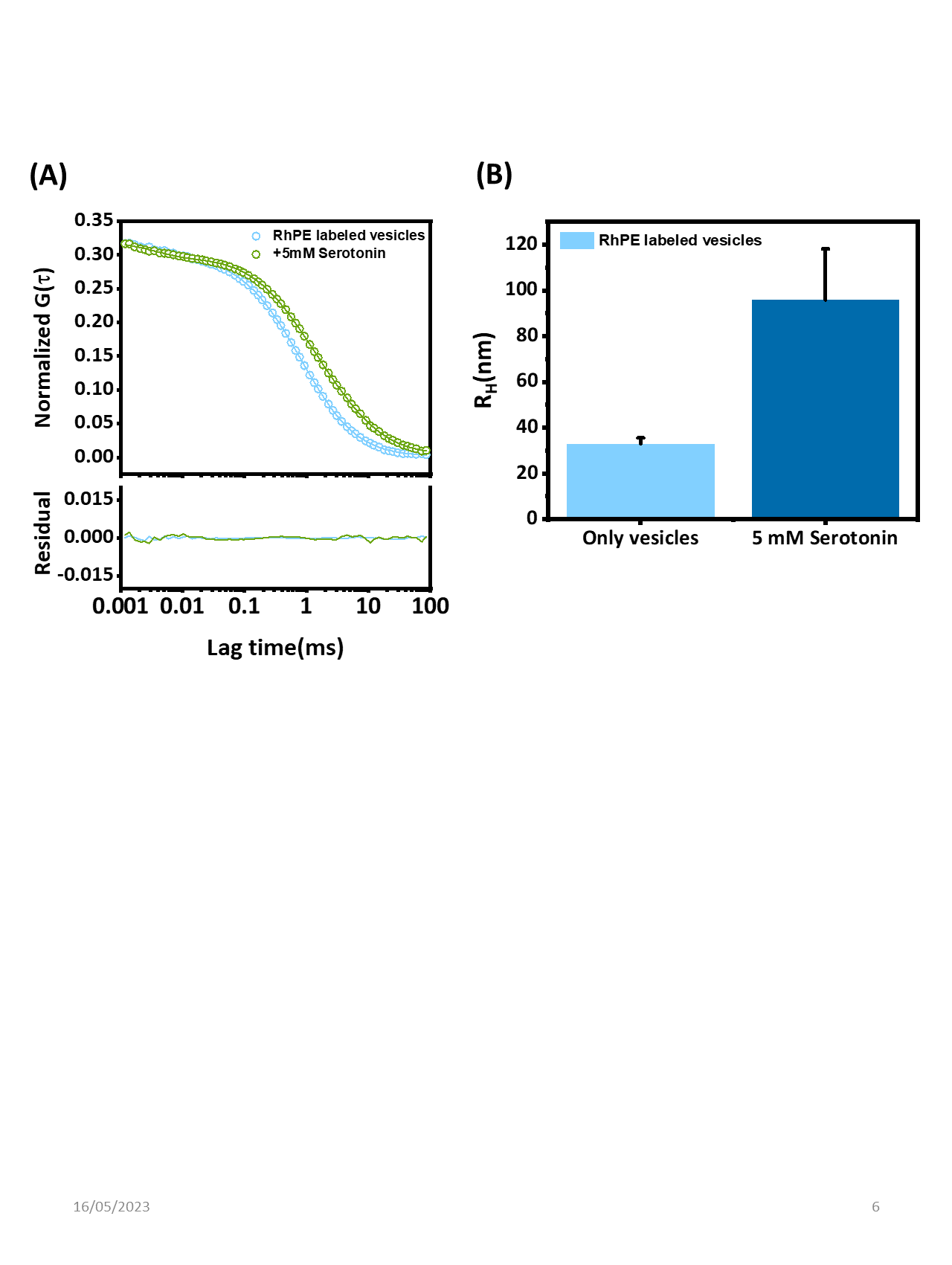

**Fig. S1: Serotonin-induced association of Rh-PE labelled PC/PE/PS/Chl 3/5/2/5 vesicles**

**(A)** Representative fluorescence autocorrelation curve of Rh-PE labelled PC/PE/PS/Chl 3/5/2/5 vesicles (data: circles) in 20 mM HEPES buffer (pH 7.4, 150 mM NaCl) with their corresponding fits (solid lines) and residuals (solid lines at the bottom). i)In the absence of serotonin (sky blue), fit: sky blue, residual: sky blue. ii)In the presence of 5 mM serotonin (dark blue), fit: dark blue, residual: dark blue. **(B)** The average hydrodynamic radius of vesicles with and without 5 mM serotonin calculated from their corresponding autocorrelation curves.

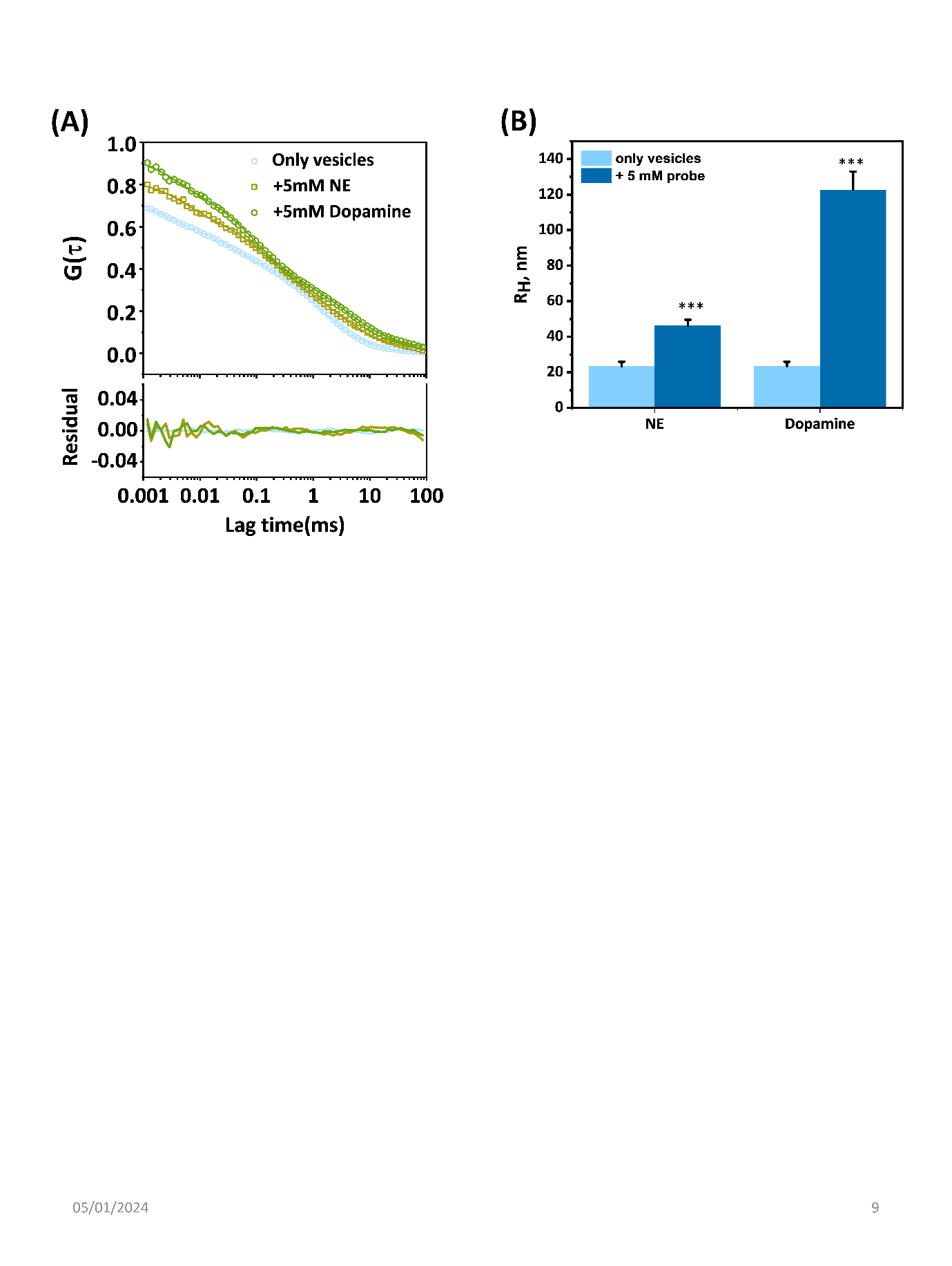

**Fig. S2: Effect of other monoamine neurotransmitters and tryptophan**. **(A)** Representative fluorescence autocorrelation curves (data: circles, squares, hexagons) with their corresponding fits (solid lines) and residuals (solid lines at the bottom) in the absence of any neurotransmitters (sky blue), in the presence of 5 mM norepinephrine (grey-green), and 5 mM dopamine (green). **(B)** Average hydrodynamic radii of vesicles in the absence and presence of (5 mM) norepinephrine, dopamine and tryptophan.

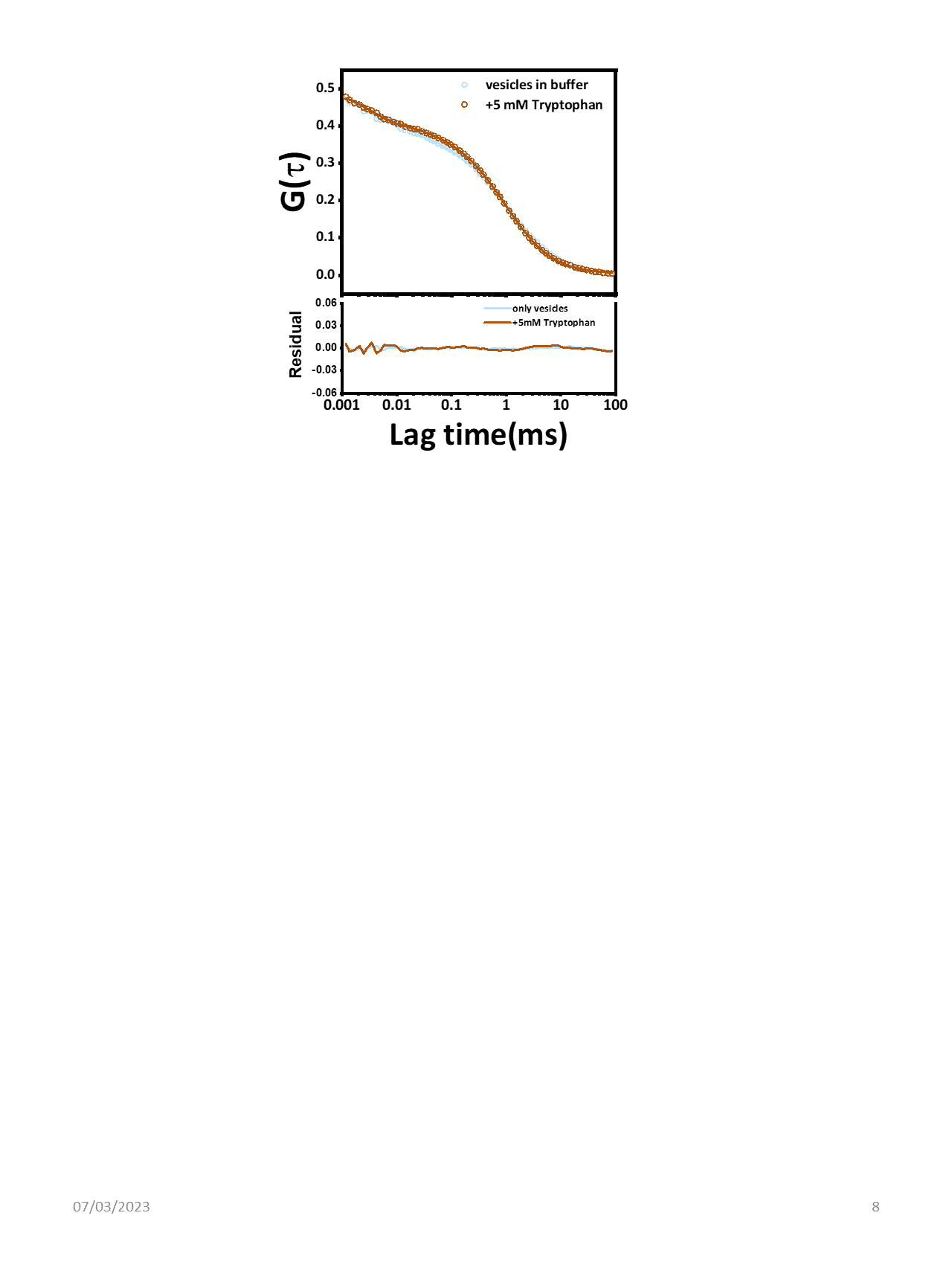

**Fig. S3:** Representative fluorescence autocorrelation curve of PC/PE/PS/Chl 3/5/2/5 vesicles fluorescently labeled with Nile Red dye i) In the absence of tryptophan (sky blue circle), fit: sky blue line, residual: sky blue line at the bottom and ii) In the presence of 5 mM tryptophan (brown circle), fit: brown line, residual: brown line at the bottom. There is no change in the decay of the autocorrelation curve in the presence of tryptophan.

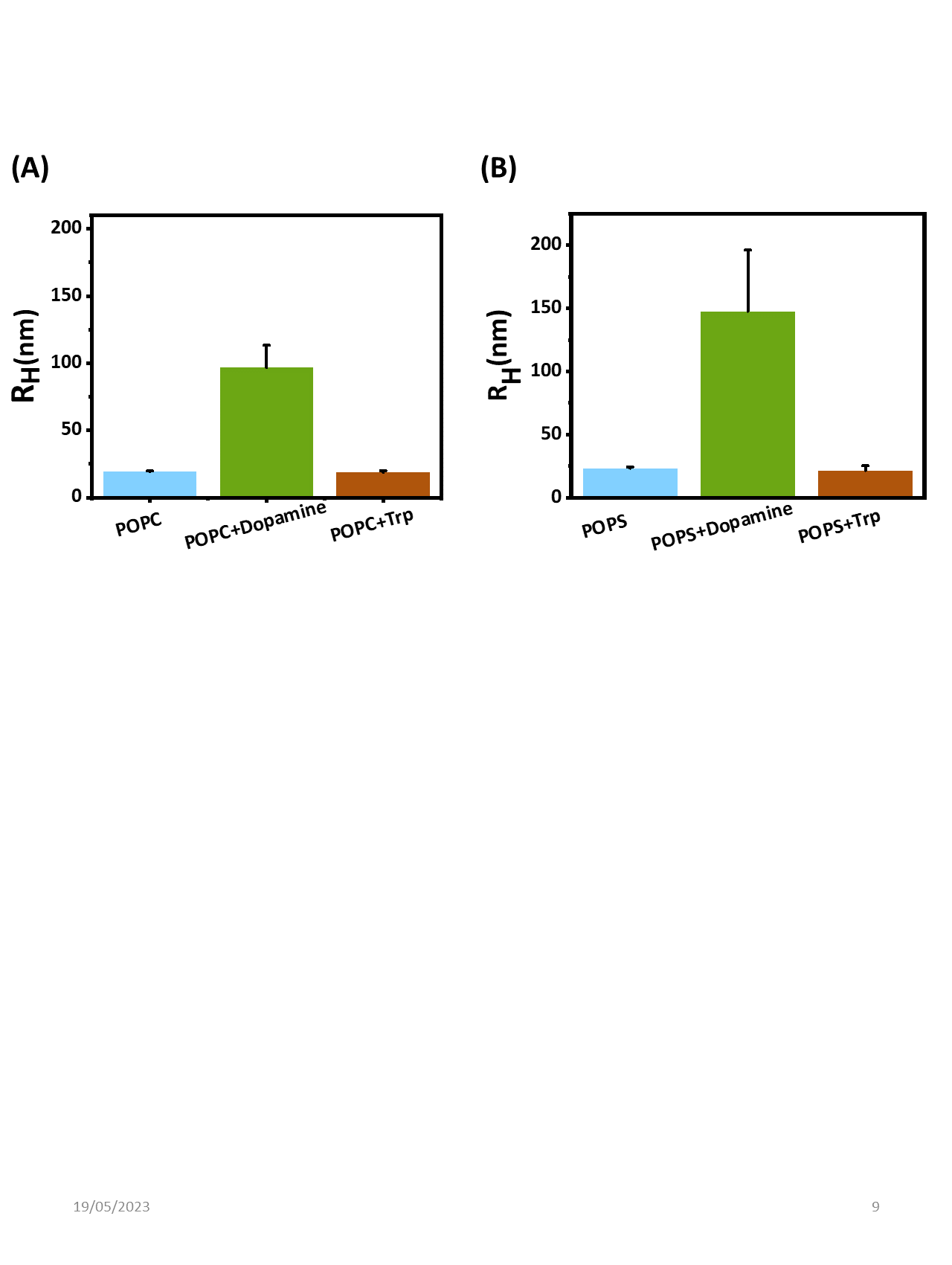

**Fig. S4:** The average hydrodynamic radius of the **(A)** POPC (neutral lipid) and **(B)** POPS (negatively charged lipid) vesicle solution in the absence (sky blue) and presence of 5 mM Dopamine (green) and 5 mM Tryptophan (brown).

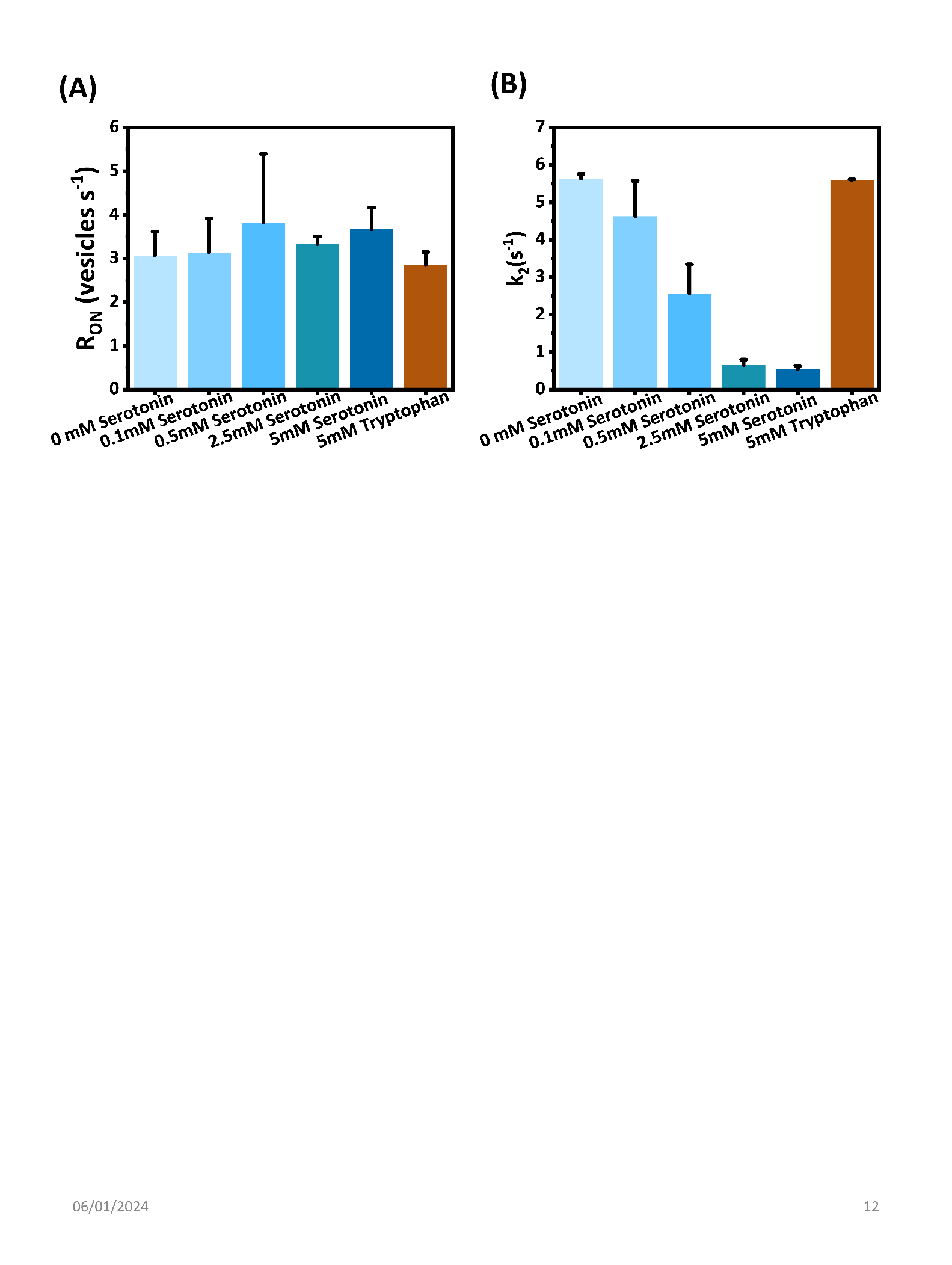

**Fig. S5:** The rate of binding (R_ON_) of vesicles with the SLBs as a function of the concentration of serotonin and in the presence of 5 mM tryptophan in the TIRF imaging experiment.

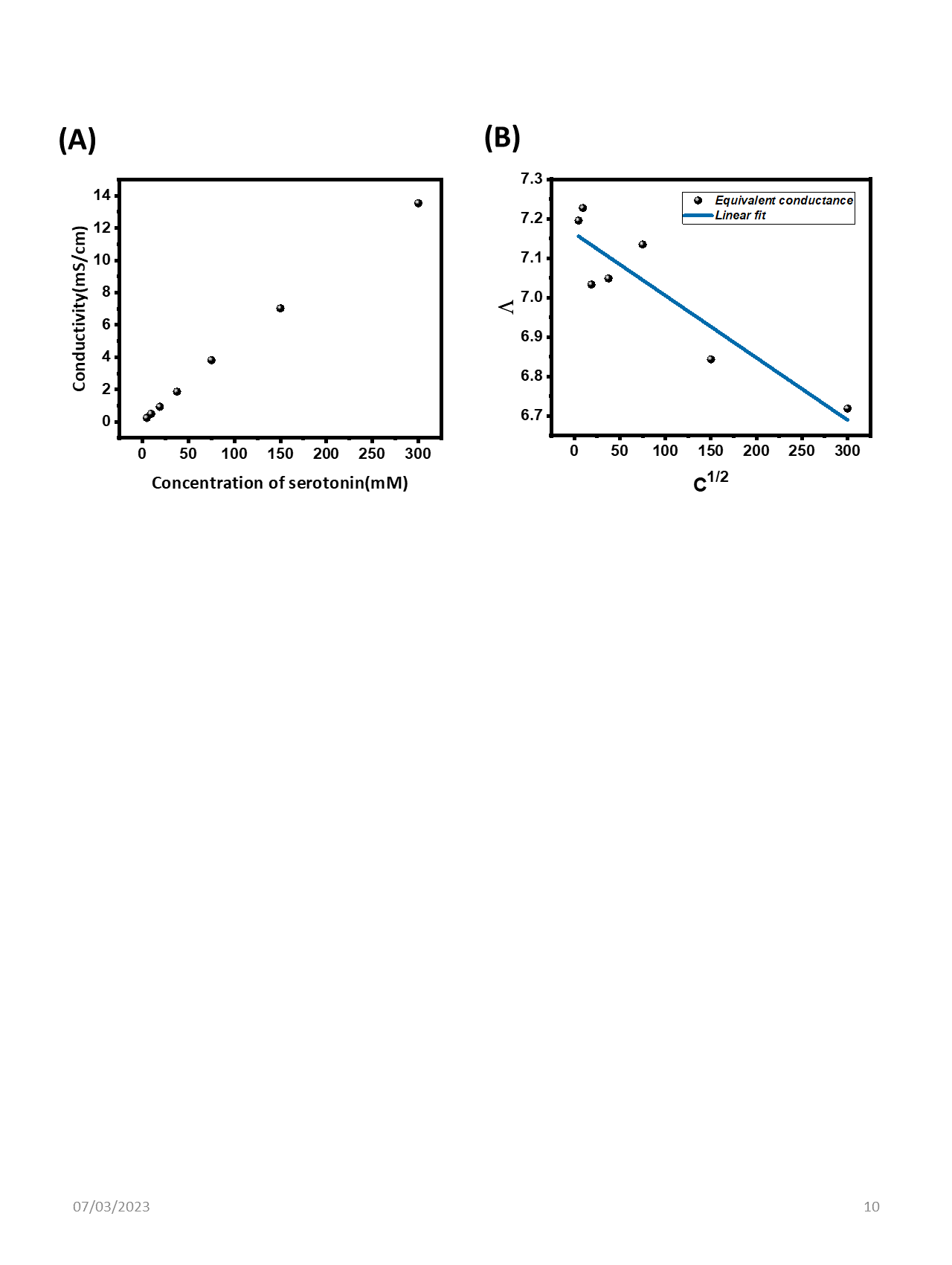

**Fig. S6:** Conductivity measurement of serotonin hydrochloride at pH 7.4. **(A)** The conductivity of the solution as a function of the concentration of serotonin hydrochloride. **(B)** The equivalent conductance of the solution calculated from the previous curve as a function of the concentration of serotonin hydrochloride. The linear nature of both curves suggests that serotonin is a strong electrolyte at pH 7.4 dissociation into two ionic species viz serotonin-H^+^ and Cl^-^.

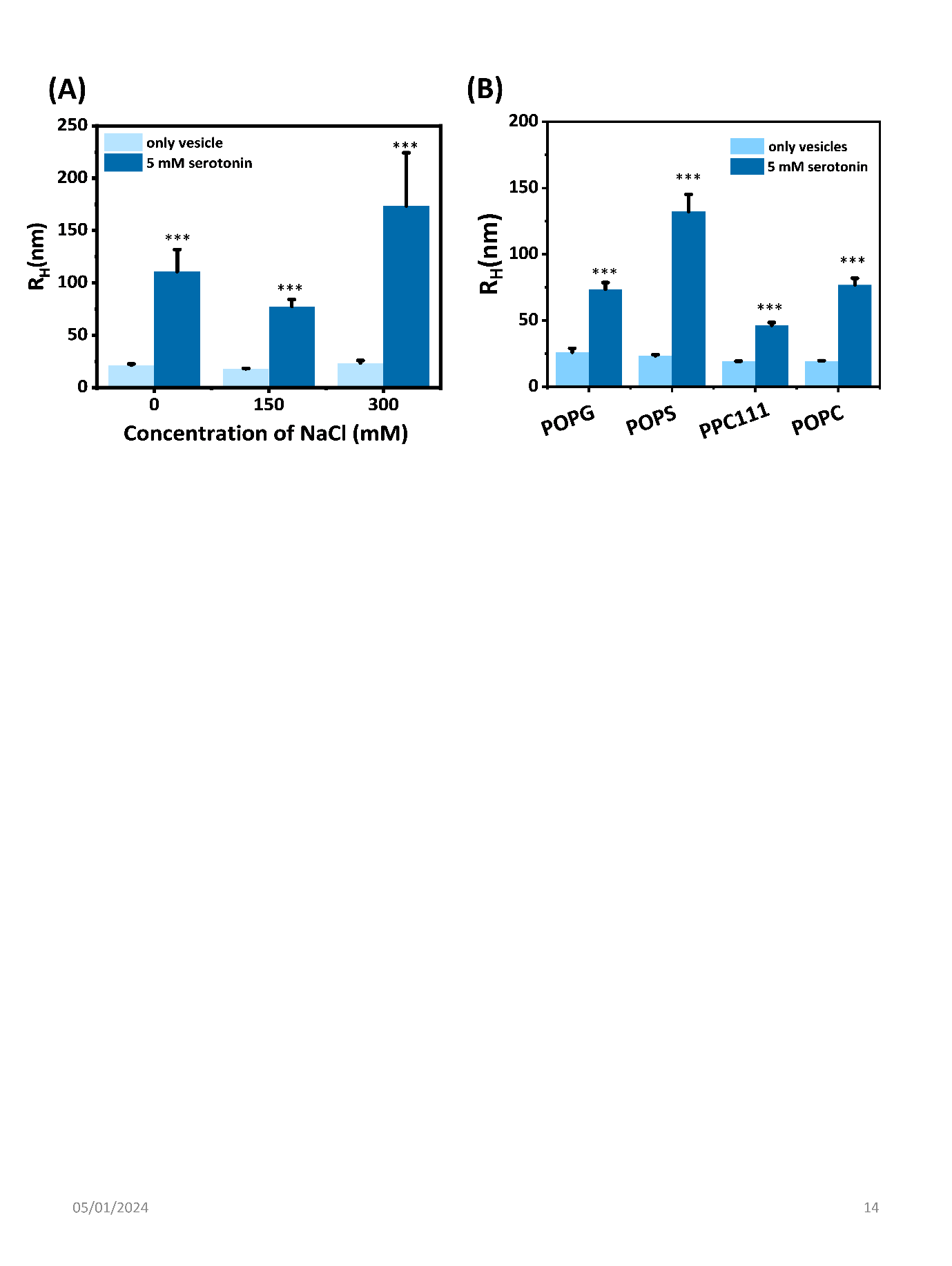

**Fig. S7: Effect of electrostatics on the serotonin-induced vesicle association as probed by FCS. (A)** Effect of concentration of NaCl (0 mM, 150 mM, 300 mM) on the serotonin-induced association behavior (R_H_) of PC/PE/PS/Chl 3/5/2/5 vesicles probed by FCS (sky blue bars: R_H_ of vesicles in buffer, dark blue bars: R_H_ of vesicles in the presence of 5 mM serotonin). **(B)** Effect of lipid membrane composition on the serotonin-induced association of vesicles probed by FCS. R_H_ of POPG, POPS, PPC111, and POPC vesicles with and without serotonin. Sky blue: vesicles in buffer, dark blue: vesicles in the presence of 5 mM serotonin.

Mean and S.E.M are plotted. This dataset passed tests for normality (Shapiro-Wilk test and Kolmogrov-Smirnov test, p>0.05) and equality of variances (F-test for variances, p>0.05). Repeated measures t-tests were performed to check the significance level. *p<0.05, **p<0.01, ***p<0.001 compared to the control (only vesicles)

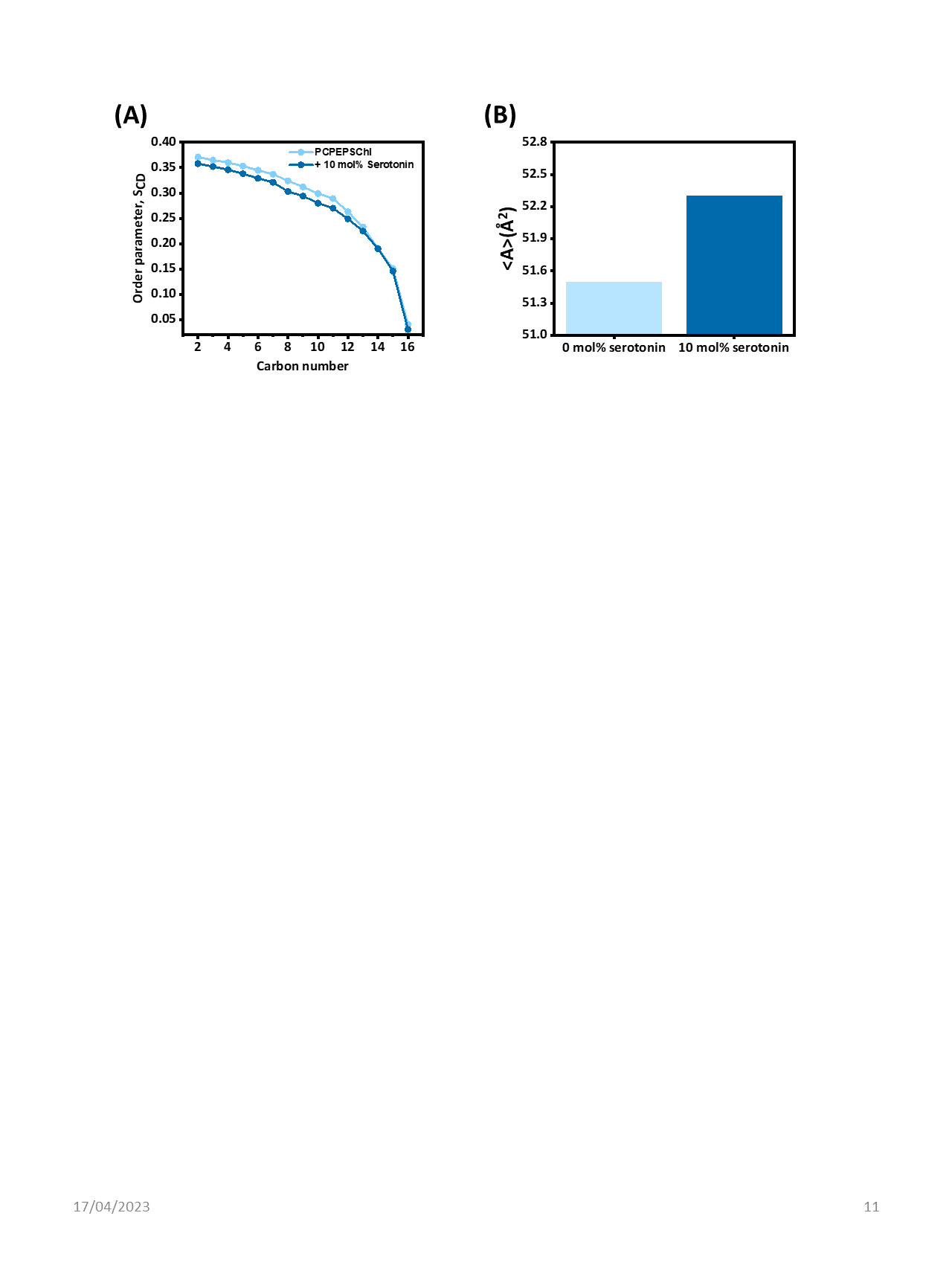

**Fig. S8: (A)** Order parameter profile along the lipid carbon chain of POPC in PC/PE/PS/Chl 3/5/2/5 membrane in the absence and presence of 10 mol% serotonin. Inset shows the average order parameter in the absence and presence of serotonin (reproduced with permission from Gupta, A.; Krupa, P.; Engberg, O.; Krupa, M.; Chaudhary, A.; Li, M. S.; Huster, D.; Maiti, S. Unusual Robustness of Neurotransmitter Vesicle Membranes against Serotonin-Induced Perturbations. *J. Phys. Chem. B* **2023**, *127* (9), 1947–1955. Copyright © 2023, American Chemical Society^10^). **(B)** The average lateral area occupied per lipid molecule of POPC in PC/PE/PS/Chl 3/5/2/5 membrane in the absence and presence of 10 mol% serotonin.

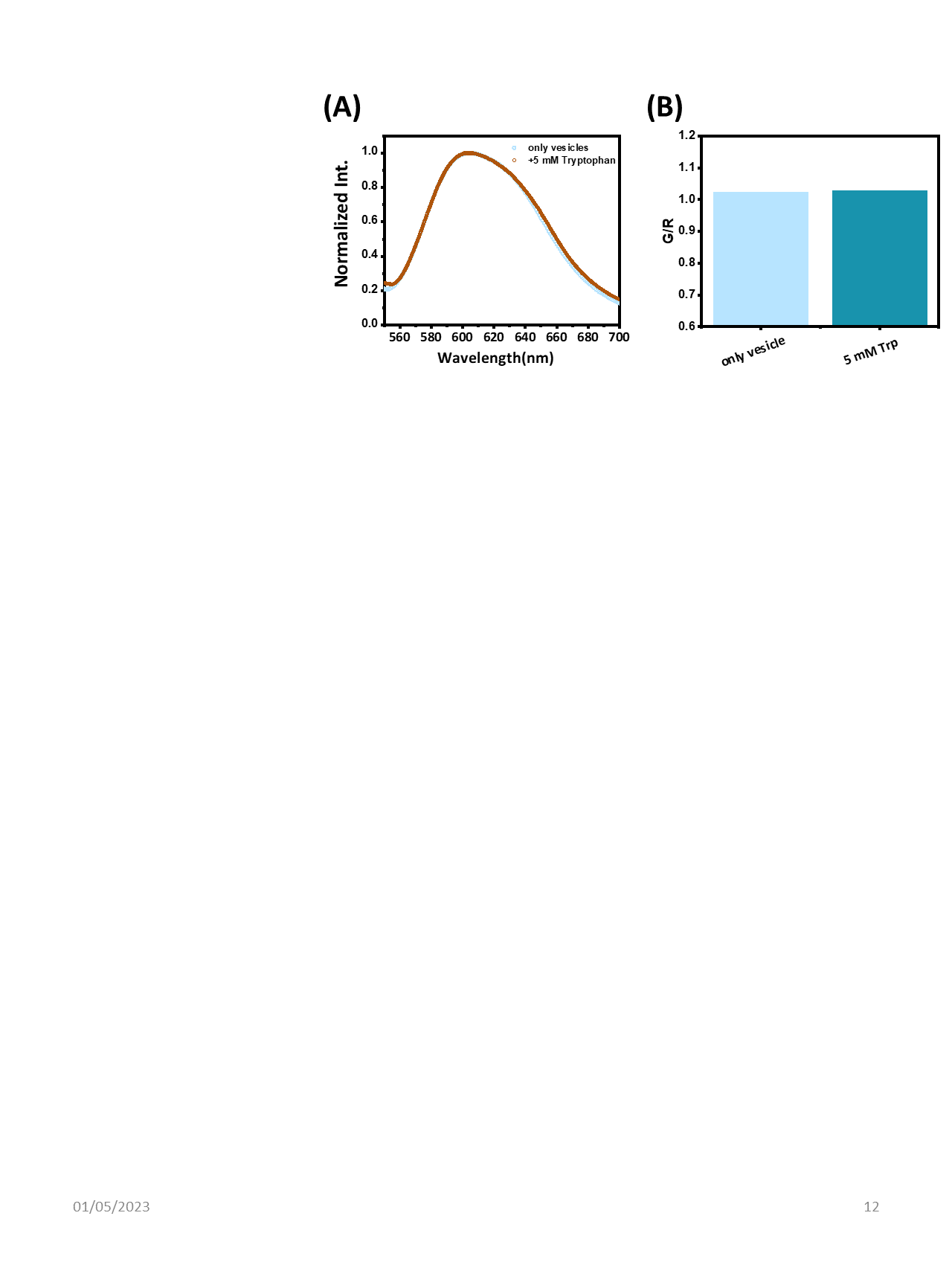

**Fig. S9: (A)** Normalized fluorescence emission spectra of Nile Red labelled PC/PE/PS/Chl 3/5/2/5 vesicle solution in the absence (sky blue) and presence (brown) of 5 mM tryptophan. **(B)** G/R ratio calculated from the Nile red spectra of vesicle solution with and without 5 mM tryptophan.

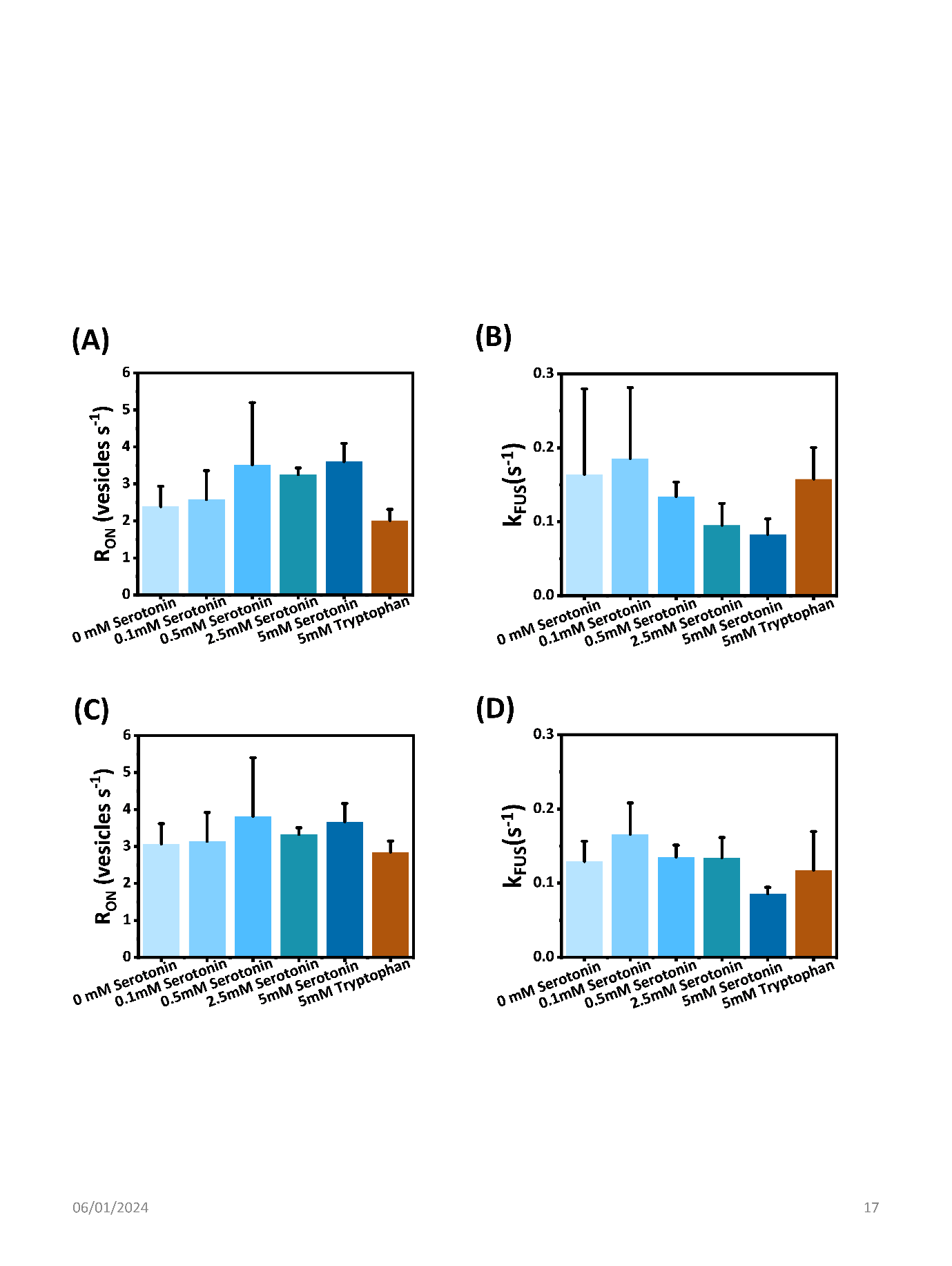

**Fig. S10:** The R_ON_ and the k_FUS_ before **(A and B)** and after **(C and D)** correction of the non-detectable events in TIRF imaging.

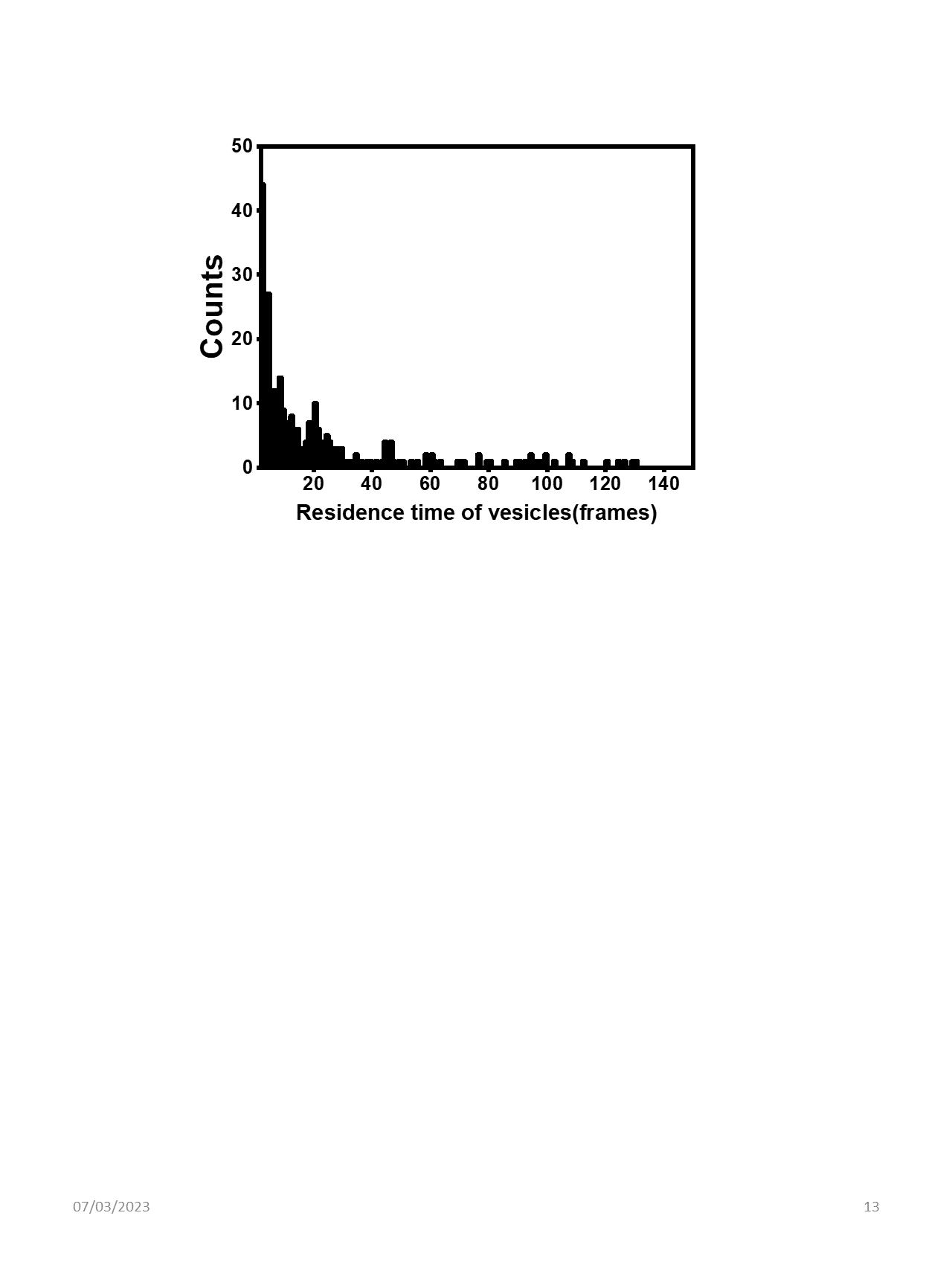

**Fig. S11:** A representative exponential distribution of residence time of vesicles in the TIRF imaging experiment
